# Determining the stability of genome-wide factors in BMI between ages 40 to 69 years

**DOI:** 10.1101/2021.07.28.454172

**Authors:** Nathan A Gillespie, Amanda Elswick Gentry, Robert M Kirkpatrick, Hermine H Maes, Chandra A Reynolds, Ravi Mathur, Kenneth S Kendler, Roseann E. Peterson, Bradley T. Webb

**Author notes:** Corresponding author: Nathan Gillespie. Shared senior authorship (Roseann E. Peterson, Bradley T. Webb).

## Abstract

Genome-wide association studies (GWAS) have successfully identified common variants associated with BMI. However, the stability of genetic variation influencing BMI from midlife and beyond is unknown. By analyzing BMI data collected from 165,717 men and 193,073 women from the UKBiobank, we performed BMI GWAS on six independent five-year age intervals between 40 and 73 years. We then applied genomic structural equation modeling (gSEM) to test competing hypotheses regarding the stability of genetic effects for BMI. LDSR genetic correlations between BMI assessed between ages 40 to 73 were all very high and ranged 0.89 to 1.00. Genomic structural equation modeling revealed that genetic variance in BMI at each age interval could not be explained by the accumulation of any age-specific genetic influences or autoregressive processes. Instead, a common set of stable genetic influences appears to underpin variation in BMI from middle to early old age in men and women alike.

## Introduction

Body mass index (BMI) is a commonly assessed trait in population studies and often used to index underweight, normal weight, overweight, or obese individuals. An individual’s BMI and how it changes can differentially predict numerous health (e.g., cardiometabolic, diabetes, dementia ^1,2^) and mortality outcomes ^3^. Despite the known limitations of BMI ^4-6^ and value added by considering additional adiposity indices such as waist-to-hip ratio and waist circumference for predicting cardio-metabolic diseases ^7^ and mortality ^8^, increased BMI still remains a significant predictor of dementia ^2,9,10^ and all-cause mortality ^11^, including more recently, COVID19-related hospitalization and death ^12^. Understanding the genetic and environmental etiology of BMI is therefore a public health priority. Among the many endeavors aimed at identifying the loci underpinning variation in complex traits ^13^, genome-wide association scan (GWAS) analyses of adult BMI ^14^ are some of the most successful. For instance, BMI meta-analytic GWAS results can now account for a significant proportion of BMI heritability ^15,16^. A limitation however, of these results is their reliance on aggregated data, which include different cohorts sampled from varying geographic and economic regions, comprising multiple birth cohorts, and age distributions. Here, we examine the last caveat because it remains to be empirically determined if genome-wide variation associated with adult BMI is age-invariant or age-specific.

### BMI heritability & longitudinal genetic correlations

Despite moderate stability across time ^1,17-20^, average adult BMI increases from age 20 to age 65 at which time it levels off until age 80 ^1^ when it begins to decline. Such changes might be attributed to variable contributions of genetic and environmental risks across the lifespan. Apart from birth cohort differences, Dahl et al. ^1^, found that factors such as an obesity genetic risk score, type-2 diabetes mellitus, cardiovascular disease, substance use, and educational attainment were all differentially predictive of both average BMI and changes in BMI before and after age 65. In contrast, many of these risks were no longer predictive after age 65.

Whereas average lifetime BMI heritability is 0.75 ^16^ with estimates range from 0.47 - 0.90 and from 0.24 - 0.81 in twin studies and family studies respectively ^16,21^ - heritability actually increases throughout infancy and adolescence ^22^ before decreasing during adulthood ^16^. Twin studies have shown that genetic influences in BMI are correlated across time from infancy to adolescence ^17-19,22^ and from early adulthood to midlife ^20,23^ and sometimes very highly ^24^, which indicates continuous expression of the same genetic influences ^25^. However, longitudinal genetic correlations for BMI never reach unity. Indeed, there is considerable variability in longitudinal genetic correlations (rg=∼.55-.95). This is consistent with age-specific genetic influences, which could be obscured in GWAS meta-analyses that model age as a covariate.

In addition to biometrical genetics or longitudinal twin designs, molecular methods such as linkage disequilibrium score regression (LDSR) ^26^ can leverage GWAS summary statistics and population reference panels to estimate correlations (rG) across unrelated and independent samples. Extending this approach to cohort studies with different ages can address the question of whether or not genetic risks in BMI are correlated across time. Currently, there is a paucity of reports examining BMI rG across time using GWAS data. Studies to date have implemented a variety of approaches and produced mixed results. For instance, Trzaskowski et al. ^27^ used LDSR to report a genetic correlation (r_g_=0.86) between BMI assessed at ages 11 and 65. Winkler et. al. ^28^ estimated Spearman rank genetic correlations between BMI assessed in populations above and below age 50, which revealed a much smaller correlations (r_g_= 0.12). Notwithstanding the need for greater precision regarding longitudinal genetic correlations, such correlations are descriptive and provide no insight regarding competing theories underlying developmental processes in BMI.

We propose that at least two theoretical mechanisms ^29^ can explain the observed continuity in genetic correlations. The first is a growth process whereby genetic or environmental factors determine the levels and rates of change in BMI over time. In this model, variances and covariances between longitudinal measures of BMI depend on individual genetic or environmental differences in growth patterns unfolding with age (“random growth curves”) ^30-34^. We are aware of three twin studies that have applied genetically informative growth models to longitudinal BMI data ^35-37^. Unfortunately, random growth curves do not determine the extent to which stability or changes in BMI are governed by time-invariant versus age-specific genetic influences. To address this question, the second mechanism predicts that variances and covariances are determined by random, time-specific genetic and environmental effects, which are more or less persistent over time i.e., autoregressive effects’ ^38-40^. Illustrated in Figure 1, this approach predicts a causal hypothesis of inertial effects, whereby genetic factors contributing to BMI at one time causally affect BMI at the next. We have applied this approach to personality ^41^, anxiety and depression ^42,43^, substance use ^44^ and brain aging ^45^. We are aware of two reports that have tested the fit of autoregession models to BMI data ^18,25^. In addition to age-invariant genetic risks, Cornes et al. ^18^ also found evidence of distinct, age-specific genetic influences on BMI at ages 12, 14 and 16. To our knowledge, autoregressive effects have not been tested in adult BMI, especially across a wide window comprising narrow age intervals in adults. Fortunately, the recent, innovative application of structural equation modelling (SEM) to LDSR genetic correlations ^46^ based on available GWAS results can now address the aforementioned gaps.

**Figure 1.**
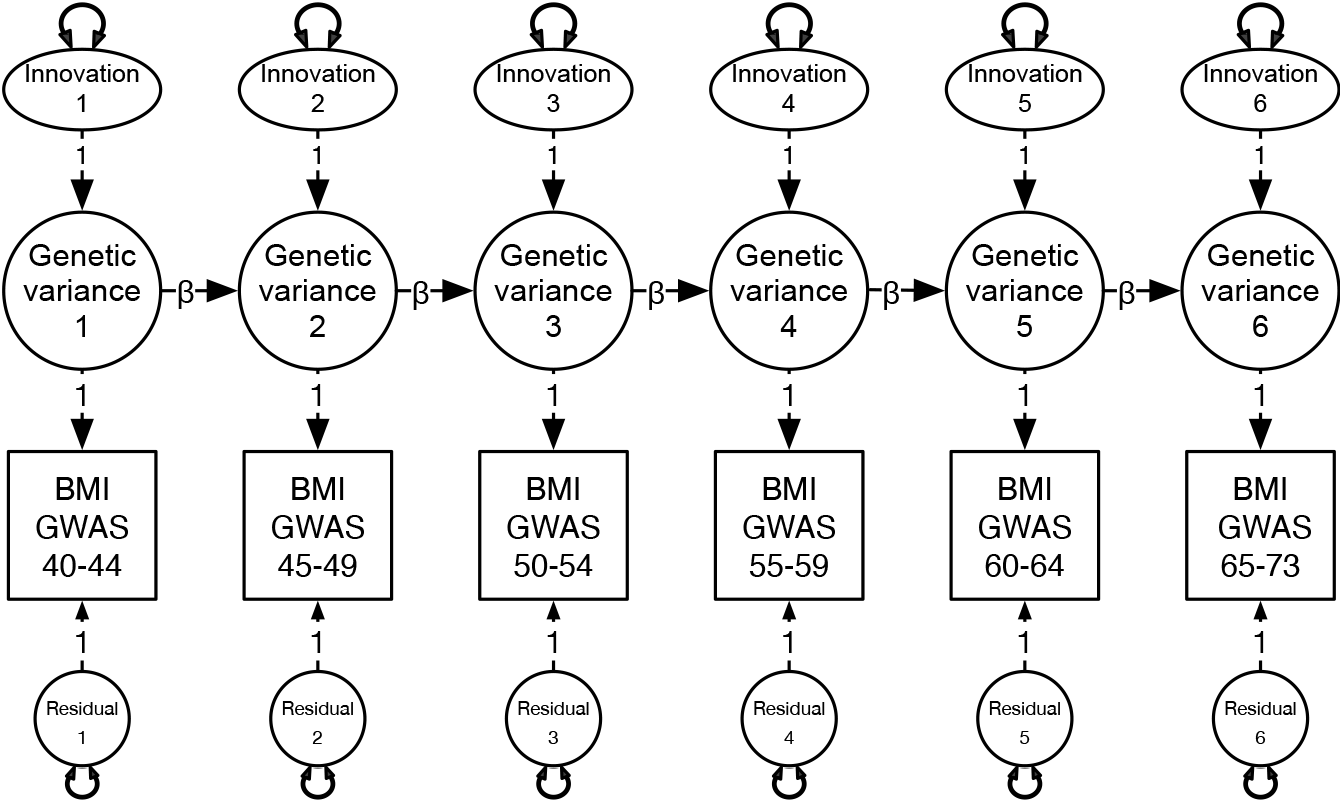
Autoregression model depicting genome-wide variation in BMI at each age interval in terms of time-specific variation or “innovations” & the causal contribution of genome-wide variation from previous age intervals. Model also includes residual genetic variation not otherwise explained by the autoregression. Note: Double-headed arrows denote variation associated with innovations & residuals at each age interval. Beta (β) denotes the causal contribution of variance from one age interval to the next.

By applying genomic SEM or “gSEM” to BMI GWAS data from the UK Biobank ^47^, our aim was to determine if genetic influences across middle age were best explained by age-dependent versus age-invariant processes. We also tested if alternative, more parsimonious theoretical explanations i.e., common factor models, whereby covariance between genetic influences across time could be captured by a single latent factor ^48^, provided a better fit to the data. Given that standardized estimates of BMI heritability for men and women are statistically equal ^16^ and that there appear to be no sex differences in terms of the observed adult decline in heritability ^49^, we hypothesized that developmental processes governing changes in heritability over time likewise ought to be comparable across sex.

## Methods

All BMI and GWAS data came from the UK Biobank, which is accessible to the scientific community, which is a large and detailed prospective study of over 500,000 subjects aged 40–73 years recruited from 2006 to 2010 of predominantly European ancestry ^47^.

### BMI data

Described in detail elsewhere ^50^, weight was collected from subjects using a Tanita BC418MA body composition analyzer. Standing and sitting height measurements were collected from subjects using a Seca 240cm height measure. Weight and height were amalgamated into mass index (BMI) calculated as weight divided by height squared (kg/m2). We divided the BMI data into six age intervals: 40-44; 45-49; 50-54; 55-59; 60-64; and 65-73 years based on age and BMI at baseline assessment. The range was based on available data whereas the number of age tranches was selected to maximize our power to choose between competing longitudinal and multivariate models without minimizing the statistical power of the GWAS analyses at each interval. The number of subjects with complete BMI and GWAS data are shown in Supplementary Table S1.

### Genotypic data

Genotype data were filtered according to the Neale Lab pipeline, using filtration parameters and scripts publicly available from the lab’s GitHub ^51^ repository. Samples were filtered to retain only unrelated subjects of British ancestry (n=359,980.) Imputed variants ^52^ were filtered based on INFO scores > 0.8, MAF > 0.001, and HWE p-value > 1e-10.

### GWAS analyses

This is proof-of-principle illustration of the application of structural equation modelling (SEM) to GWAS summary data to test longitudinal hypotheses. While a subset of UKB subjects has repeated BMI measures, the data freeze used here contained insufficient subjects with a minimum of three BMI assessments required to fit autoregression models. To maximize power and subject inclusion, we divided the sample into 5-year age intervals, which yielded 6 age tranches for GWAS and the subsequent gSEM pseudo-longitudinal analyses. Separate GWAS analyses were conducted for each 5-year age interval using BGENIE (version 1.3.) ^52^. The first 10 ancestry principal components were included as covariates in all models and sex was also included as a covariate in the non-sex stratified models.

### Genomic structural equation modelling

We then applied the gSEM software package ^46^ in R (version 4.0.3) ^53^ to estimate the genetic variance-covariance (S) and asymptotic sampling covariance ‘weight’ (V) matrices based on the 6 BMI GWAS results combined across, and second, based on the separate GWAS results by sex. Estimation of the S and V matrices is a 3-step process. In step 1, the raw GWAS summary statistics were manipulated using the gSEM *munge* option to remove all SNPs with MAF < 1%, information scores < 0.9, and SNPs in the MHC region. In step 2, we used the gSEM *ldsc* option to run multivariate LD score regression ^46^ to estimate the covariance (S) and asymptotically weighted covariance (V) matrices between the GWAS summary statistics. In step 3, we then used the lavaan (version 0.6-7) ^54^ in R (version 4.0.3) ^53^ to fit and compare competing longitudinal and multivariate models to the S and V matrices.

The autoregression model predicts that time-specific random genetic or environmental effects “innovations” are more or less persistent over time (autoregressive effects) ^38^. As described by Eaves and others ^38-40^ and illustrated in Figure 1, genetic variance at each occasion is a function of i) new random effects “innovations” on the phenotype as well as ii) the linear contribution of genetic differences expressed at the preceding time. We assume that cross-temporal correlations arise because the innovations have a more or less persistent effect over time and may, under some circumstances accumulate, potentially giving rise to developmental increases in genetic variance and increased correlations between adjacent measures. One consequence of the autoregressive model is the tendency of cross-temporal correlations to decay as a function of increasing lag-time. Depending on the magnitude of an innovation and its relative persistence, the observed variances and cross-temporal covariances may increase towards a stable asymptotic value. We tested this autoregression model by fitting genetic innovations at the 45-49, 50-54, 55-59, 60-64 and 65-73 age-intervals, which we then successively dropped. We then specified an autoregression model that included a single source of genetic variance at the 40-44 age interval, which accounted for genetic variance at all subsequent age intervals. Finally, we fitted an exploratory factor analysis comprising a single factor.

### Model fit indices & comparisons

In gSEM analyses there is no one sample size to speak of. This is because GWAS studies from which the summary statistics are derived can vary in size and subject overlap. Thus, potentially, a different (effective) sample size may apply to each element of S. We were therefore limited to fit indices that do not explicitly depend upon sample size: the pseudo Akaike Information Criterion (pseudoAIC); Comparative Fit Index (CFI); Tucker Lewis Index (TFI); and the Standardized Root Mean Square Residual (SRMR) to judge the best-fitting model. Both the CFI and TFI are incremental fit indices that penalize models with increasing complexity. The SRMR is an absolute measure of fit based on the difference between the observed and predicted correlations under each model, such that a value of zero indicates a perfect fit. The pseudoAIC is a comparative fit index, whereby the model with the lowest AIC values is interpreted as the best-fitting.

## Results

### Combined male & female analyses

The LDSR-based genome-wide genetic correlations between the six GWAS summary statistics, including GWAS sample sizes and SNP-based heritability at each age interval, are shown in Table 1. The correlations do not continue to decline with increasing time intervals, which would be indicative of a simplex structure best explained by autoregression models. For example, the LDSR genetic correlation (r_g_) between BMI at ages 40-45 and 66-73 years was higher than the r_g_ between BMI at ages 40-45 and 56-60 years (r_g_=0.97 vs 0.93). Overall, the genetic correlations were very high and ranged from 0.93 to 1.00.

**Table 1.**
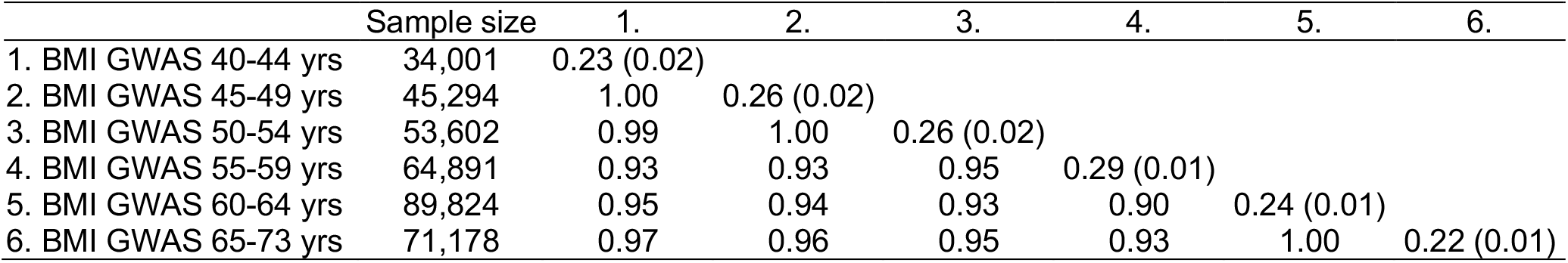
Sample sizes, estimates of SNP-based heritability (including standard errors along diagonal) & linkage disequilibrium score regression genetic correlations between the six age intervals based on the combined male & female GWAS summary statistics.

|  | Sample size | 1. | 2. | 3. | 4. | 5. | 6. |
| --- | --- | --- | --- | --- | --- | --- | --- |
| 1. BMI GWAS 40-44 yrs | 34,001 | 0.23 (0.02) |  |  |  |  |  |
| 2. BMI GWAS 45-49 yrs | 45,294 | 1.00 | 0.26 (0.02) |  |  |  |  |
| 3. BMI GWAS 50-54 yrs | 53,602 | 0.99 | 1.00 | 0.26 (0.02) |  |  |  |
| 4. BMI GWAS 55-59 yrs | 64,891 | 0.93 | 0.93 | 0.95 | 0.29 (0.01) |  |  |
| 5. BMI GWAS 60-64 yrs | 89,824 | 0.95 | 0.94 | 0.93 | 0.90 | 0.24 (0.01) |  |
| 6. BMI GWAS 65-73 yrs | 71,178 | 0.97 | 0.96 | 0.95 | 0.93 | 1.00 | 0.22 (0.01) |

Formal model fitting comparisons are shown in Table 2. We began with a fully saturated autoregression model comprising unique genetic influences or innovations at each age interval (Figure 1). This provided a reasonable fit to the data as judged by the non-significant chi-square, very high CFI and TLI values and very low SRMR. Autoregression sub-models in which the genetic innovations at ages 65-73, 60-64, and 65-73, 60-64, 55-59, 50-54 and 45-49 years were each successively removed provided only marginal improvements in terms of their pseudoAIC values. In contrast, the model with a single factor provided the overall best fit in terms of the smallest chi-square, lowest pseudoAIC and lowest SRMR. Under this model (see Figure 2), genetic variance at each five-year age interval was best explained by a single factor with a genome-wide SNP heritability of 24%.

**Table 2.** Multivariate modeling fitting comparisons based on the combined male & female GWAS summary statistics.

| Models | Chi-square(df) | p | pseudoAIC | CFI | TLI | SRMR |
| --- | --- | --- | --- | --- | --- | --- |
| Full auto-regression (AutoReg) | 21.113<sub>(13)</sub> | 0.071 | 55.113 | 0.999 | 0.999 | 0.039 |
| AutoReg: genetic innovation at 65-73 yrs dropped | 22.419<sub>(14)</sub> | 0.070 | 54.419 | 1.000 | 0.999 | 0.039 |
| AutoReg: genetic innovation at 60-64 yrs dropped | 20.872<sub>(14)</sub> | 0.105 | 52.872 | 1.000 | 0.999 | 0.039 |
| AutoReg: genetic innovation at 55-59 yrs dropped | 25.403<sub>(14)</sub> | 0.031 | 57.403 | 0.999 | 0.999 | 0.041 |
| AutoReg: genetic innovation at 50-54 yrs dropped | 21.768<sub>(14)</sub> | 0.084 | 53.768 | 0.999 | 0.999 | 0.040 |
| AutoReg: genetic innovation at 45-49 yrs dropped | 34.073<sub>(14)</sub> | 0.002 | 66.073 | 0.998 | 0.998 | 0.051 |
| AutoReg: genetic innovations at 45-73 yrs dropped | 46.133<sub>(14)</sub> | 0.000 | 70.133 | 0.998 | 0.998 | 0.056 |
| Exploratory factor analysis - 1 factor | 13.005<sub>(9)</sub> | 0.162 | 37.005 | 1.000 | 1.000 | 0.016 |

Note: AIC = Akaike Information Criterion, CFI = comparative fit indice, TLI = Tucker Lewis Indice, SRMR = (Standardized) Root Mean Square Residual

**Figure 2.**
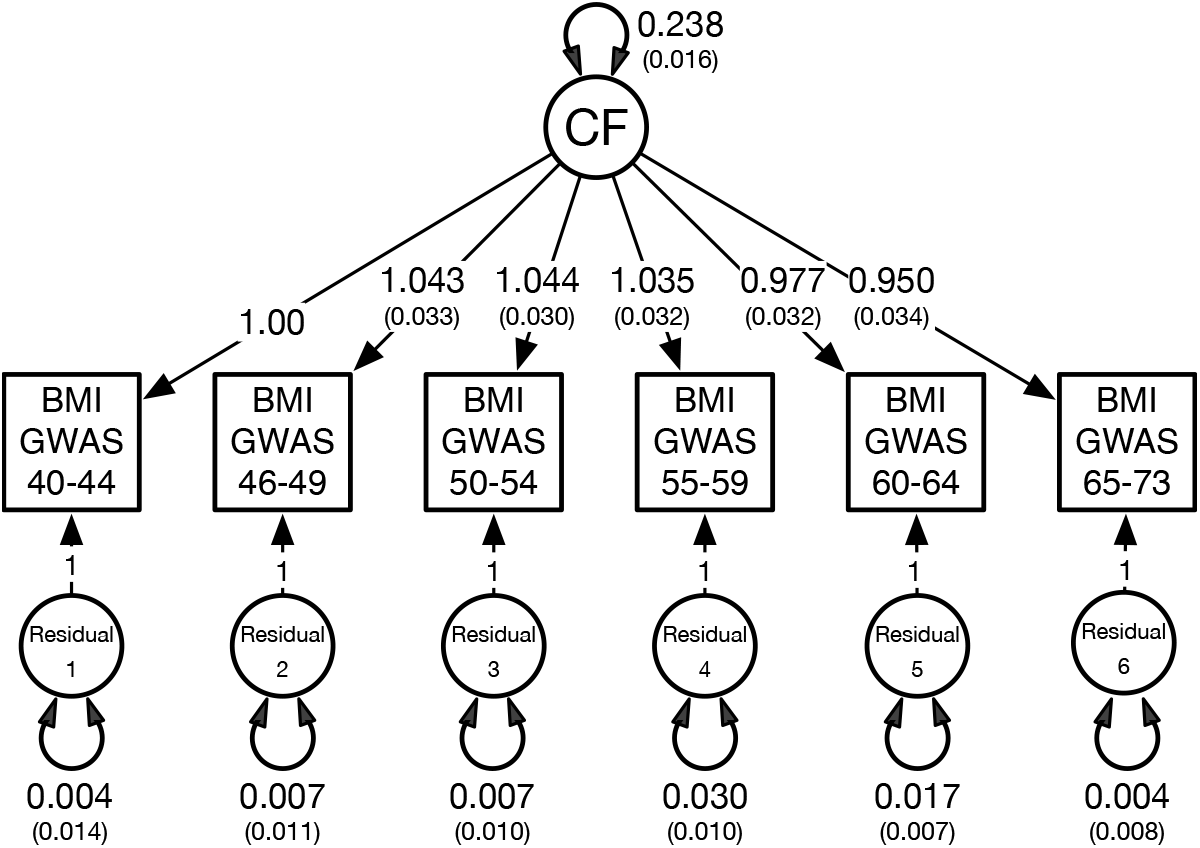
Best-fitting common factor (CF) model based on the combined male and female GWAS data. This model best explains the covariation between the six GWAS summary statistics based on five-year age intervals between ages 40-73 years. To identify this model, the factor loading on the BMI GWAS at 40-45 years was constrained to one. The double-headed arrow on the CF denotes the standardized variance, or SNP-based heritability, for BMI. Double-headed arrows on the residuals denote genetic variation at each age interval not otherwise explained by the CF.

### Sex specific analyses

An identical pattern emerged when the model fitting was repeated by sex. Male and female sample sizes at each age interval are shown in Supplementary Table S1. Table 3 shows the LDSR genetic correlations for men and women. Varying only slightly, the separate male and female genetic correlations were again high and ranged from r_g_=0.88 to r_g_=1.00. Supplementary Table S1 also shows the SNP-based heritability estimates by sex, which were not only very similar at each age interval, but were largely unchanged when 40 ancestry principal components were included as covariates.

**Table 3.** Linkage disequilibrium score regression genetic correlations based on the male (below diagonal) & female (above diagonal italics) GWAS summary statistics at six age intervals.

|  | 1. | 2. | 3. | 4. | 5. | 6. |
| --- | --- | --- | --- | --- | --- | --- |
| 1. BMI GWAS 40-44 yrs | 1 | 0.99 | 0.99 | 0.91 | 0.95 | 0.92 |
| 2. BMI GWAS 45-49 yrs | 0.98 | 1 | 1.00 | 0.96 | 0.93 | 0.93 |
| 3. BMI GWAS 50-54 yrs | 1.00 | 0.99 | 1 | 0.95 | 0.91 | 0.90 |
| 4. BMI GWAS 55-59 yrs | 0.93 | 0.89 | 0.93 | 1 | 0.88 | 0.93 |
| 5. BMI GWAS 60-64 yrs | 0.97 | 0.90 | 0.95 | 0.88 | 1 | 0.98 |
| 6. BMI GWAS 65-73 yrs | 0.97 | 0.90 | 0.96 | 0.93 | 0.99 | 1 |

As shown in Table 4, the genetic innovation parameters at ages 46-73 years for each sex could each be dropped from the full autoregression model as judged by the non-significant chi-square value. Overall, however, a model with a single common factor again provided the best fit to the data for each sex in terms of the lowest chi-square, pseudoAIC and SRMR values (see Figure 3). This suggests that there is no evidence of age-specific genetic variation in BMI for either men or women.

**Table 4.** Multivariate modeling fitting comparisons based on the combined MALE GWAS summary statistics at six age intervals.

| <u>Women</u> | ChiSquare<sub>df</sub> | p | pseudoAIC | CFI | TLI | SRMR |
| --- | --- | --- | --- | --- | --- | --- |
| Full auto-regression (AutoReg) | 15.019<sub>(13)</sub> | 0.306 | 49.019 | 1.000 | 1.000 | 0.043 |
| AutoReg: genetic innovation at 65-73 yrs dropped | 14.866<sub>(14)</sub> | 0.387 | 46.866 | 1.000 | 1.000 | 0.043 |
| AutoReg: genetic innovation at 60-64 yrs dropped | 14.883<sub>(14)</sub> | 0.386 | 46.883 | 1.000 | 1.000 | 0.043 |
| AutoReg: genetic innovation at 55-59 yrs dropped | 16.813<sub>(14)</sub> | 0.266 | 48.813 | 0.999 | 0.999 | 0.046 |
| AutoReg: genetic innovation at 50-54 yrs dropped | 14.213<sub>(14)</sub> | 0.434 | 46.213 | 1.000 | 1.000 | 0.043 |
| AutoReg: genetic innovation at 45-49 yrs dropped | 21.482<sub>(14)</sub> | 0.090 | 53.482 | 0.998 | 0.998 | 0.057 |
| AutoReg: genetic innovations at 45-73 yrs dropped | 25.617<sub>(14)</sub> | 0.109 | 49.617 | 0.998 | 0.999 | 0.059 |
| Exploratory factor analysis - 1 factor | 8.832<sub>(9)</sub> | 0.453 | 32.832 | 1.000 | 1.000 | 0.023 |
| <u>Men</u> | ChiSquare<sub>df</sub> | p | pseudoAIC | CFI | TLI | SRMR |
| Full auto-regression (AutoReg) | 11.858<sub>(13)</sub> | 0.539 | 45.858 | 1.000 | 1.000 | 0.054 |
| AutoReg: genetic innovation at 65-73 yrs dropped | 12.814<sub>(14)</sub> | 0.541 | 44.814 | 1.000 | 1.000 | 0.054 |
| AutoReg: genetic innovation at 60-64 yrs dropped | 12.085<sub>(14)</sub> | 0.599 | 44.085 | 1.000 | 1.000 | 0.053 |
| AutoReg: genetic innovation at 55-59 yrs dropped | 11.889<sub>(14)</sub> | 0.615 | 43.889 | 1.000 | 1.000 | 0.053 |
| AutoReg: genetic innovation at 50-54 yrs dropped | 21.826<sub>(14)</sub> | 0.082 | 53.826 | 0.998 | 0.998 | 0.064 |
| AutoReg: genetic innovation at 45-49 yrs dropped | 12.718<sub>(14)</sub> | 0.549 | 44.718 | 1.000 | 1.000 | 0.059 |
| AutoReg: genetic innovations at 45-73 yrs dropped | 1628.983<sub>(14)</sub> | 0.000 | 1652.983 | 0.168 | 0.001 | 0.803 |
| Exploratory factor analysis - 1 factor | 4.398<sub>(9)</sub> | 0.883 | 28.398 | 1.001 | 1.000 | 0.018 |

Note: AIC = Akaike Information Criterion, CFI = comparative fit indice, TLI = Tucker Lewis Indice, SRMR = (Standardized) Root Mean Square Residual

**Figure 3.**
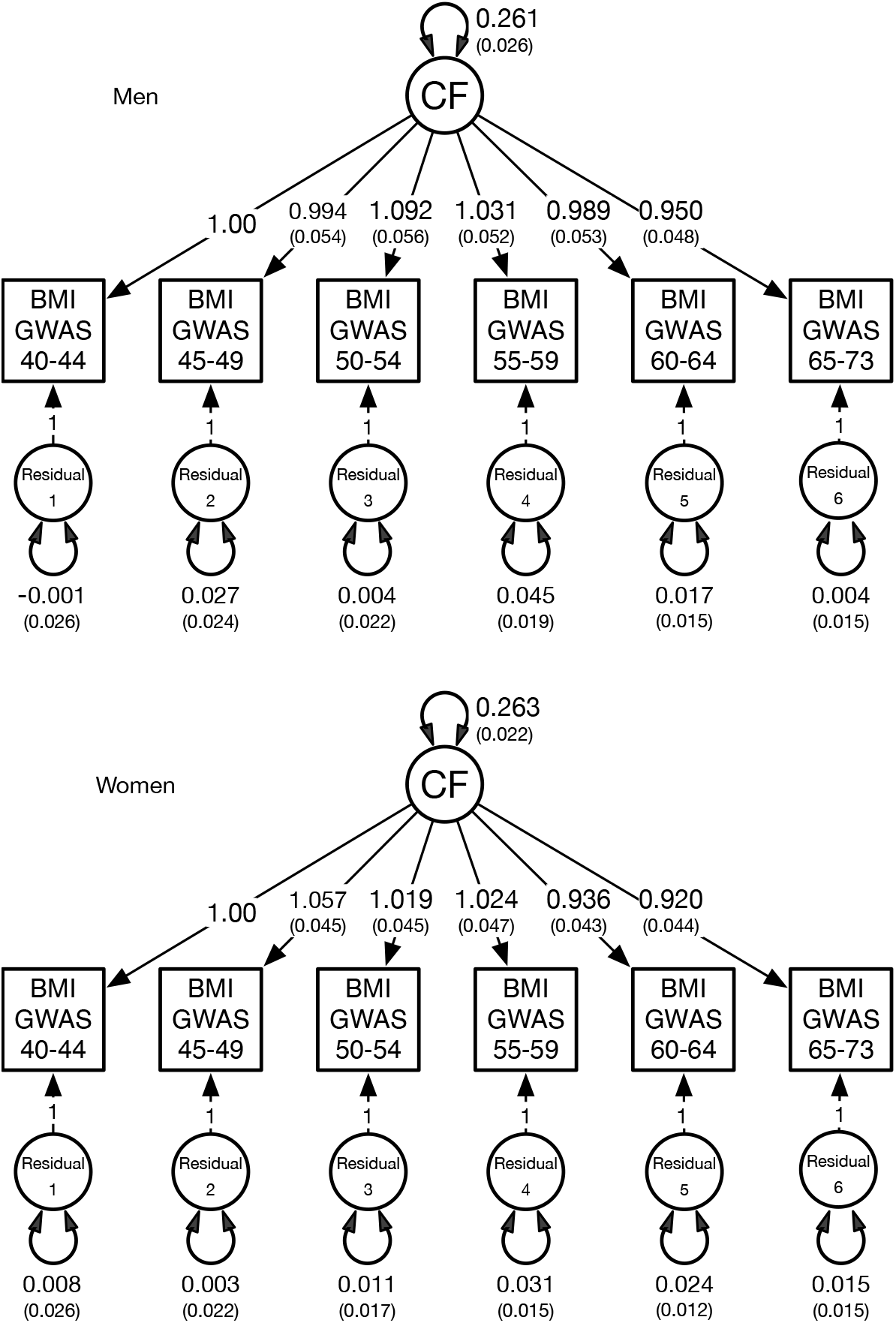
Best-fitting common factor (CF) models based on separate male and female GWAS data. To identify these models, the factor loadings on the BMI GWAS at 40-45 years were constrained to one. The double-headed arrow on the CF denotes the standardized variance, or SNP-based heritability, for BMI. Double-headed arrows on the residuals denote genetic variation at each age interval not otherwise explained by the CF.

## Discussion

This is the first study to test a genetic developmental theory of BMI heritability using molecular data and structural equation modeling. From ages 40 and 73, changes in BMI heritability could not be explained by age-specific genetic factors or an accumulation of new sources of genetic variance over time. Instead, individual differences in BMI genetic variance across this time span were best explained by a single or common set of stable, genetic influences that are observable in early midlife. This pattern was observed in both men and women.

Dahl et al.’s ^1^ analysis of Swedish twin data revealed that for men and women, BMI increases across midlife, before leveling off at 65 years and declining at approximately age 80. The extent to which the inflexion at age 65 is characterized by genetic innovations was unsupported by our results. In fact, we found that the genetic correlation between the GWAS based on BMI assessed at ages 60-64 and the other age intervals were all equally high. Thus, the genetic variance at age 60-64 does not appear to be linked to any age-specific or distinct genetic processes occurring around this time.

The extent to which our observed pattern of homogenous and stable genetic influences is replicable at younger ages remains to be determined. Longitudinal genetic correlations assessed at intervals spanning infancy, adolescence and young adult range from moderate to high ^17,19,55,56^. Felix et al. ^57^ estimated a high genetic correlation between childhood and adult BMI of 0.73. Typically, however, higher correlations are observed between shorter intervals. For example, Silventoinen et al. ^17^ reported an r_g_ of 0.31 between BMI at ages 1 and 18 that increased to 0.93 between BMI at ages 17 and 18 years. Haberstick et al. ^19^ reported a similarly high genetic correlation (r_g_ = 0.97) between BMI at ages 17 and 22 years. The autoregressive modeling reported by Cornes et al. ^18^ found evidence of distinct, age-specific genetic influences on BMI at ages 12, 14 and 16. However, the authors also found large transmission coefficients in both sexes, which means that same genetic factors were largely responsible for the observed variation across the three measurement occasions. In older samples, longitudinal genetic correlations between BMI assessed at age 20 and when subjects were in their mid-forties range from 0.60 to 0.69 ^20,23^. In terms of GWAS findings, many adult BMI variants have also been observed in early growth traits in childhood, but not infancy ^58^. We note that Sovio et al. identified one the clearest examples using molecular data of age-specific genetic effects, whereby the effect of the FTO locus on BMI reverses between adiposity peak adiposity rebound during infancy ^59^. In terms of LDSR findings, Trzaskowski et al. ^27^ reported a genetic correlation of 0.86 between BMI at ages 11 and 65. The same authors also found that the adult PRS for BMI explained at most 10% of the phenotypic variance in childhood BMI. Combined, the moderate to high longitudinal genetic correlations based on twin and molecular data suggest that individual differences in heritability spanning infancy, adolescence and early adult are likely to be explained by a combination of mostly age-invariant, and age-specific genetic influences.

### Limitations

Our findings should be interpreted in the context of four limitations.

First, the BMI data were not based on repeated measures. Consequently, our approach relied on between-subject analyses that assumed no year of birth or cohort effects. Analyses of Danish and Swedish twin data have shown that increases in mean BMI in successively younger cohorts have been accompanied by larger standardized estimates of heritability ^60,61^. To examine if cohort effects exist, we inspected the LDSR genetic correlations between the youngest and oldest age intervals i.e., two maximally age-discrepant samples of unrelated individuals. The genetic correlation was r_g_ = 0.97 (see Table 1), which suggests that the impact of cohort-related genetic heterogeneity, if present, was likely to be minimal.

Second, the UKB recruitment process did not represent a random sample of the UK population ^62^. Subjects were predominately European ancestry, more likely to be older, female, to live in less socioeconomically deprived areas than nonparticipants, and when compared with the general population, were also less likely to be obese, to smoke, and to drink alcohol daily while reporting fewer self-reported health conditions ^63,64^. Although Silventoinen et al.’s meta-analysis of twin data reported only minor differences in BMI heritability across divergent cultural-geographic regions ^65,66^, the extent to which the molecular-based genetic covariance structure observed here generalizes to non-European populations remains to be determined.

Third, while our results illustrate the flexibility of SEM in terms of its application to GWAS data to test a theory of longitudinal change, our modeling was not exhaustive. For instance, we were unable to test the hypothesis that changes in heritability could be better explained either by latent growth processes or mixture distributions ^67,68^ capable of discerning distinct genetic trajectories. We emphasize that the current method is limited to the analysis of summary variance-covariance matrices derived from the GWAS analysis of common variants. GenomicSEM does not model observed phenotypic information. Consequently, sample means could not be generated with which to model latent growth or mixture distributions. We also did not test hypotheses regarding sex differences other than to report results by sex. Dubois’ meta-analysis of 23 twin birth-cohorts found evidence of sex-limitation in terms of greater genetic variance in boys in early infancy through to 19 years ^69^. In contrast, Elks et al.’s meta-regression of 88 twin-bases estimates of BMI heritability found no evidence of sex effects ^16^. It remains to be determined if the observed minor differences in the genetic covariances and the ultimate, best fitting single-factor structure are empirically equivalent across sex.

Finally, our genomic modelling was based on aggregated GWAS summary data and so was entirely independent of environmental risks, which are known to be significant in the etiology of complex traits ^70^. Consequently, our current approach precludes modeling the contribution of environmental influences with increasing age ^71^ or making allowances for phenomenon such as genetic control of sensitivity to the environment i.e., GxE interaction ^65^, which may explain the previously observed decreases in heritability during adulthood ^16^. In this regard, methods capable of simultaneously modelling the joint effect of genes and environment are likely to prove more informative. For instance, innovative approaches capable of applying genomic-relatedness based restricted maximum-likelihood ^72^ to structural equation modeling software packages such as OpenMx ^73^ have the potential to analyze individual GWAS and phenotypic data and hold promise.

### Conclusion

In absence of true longitudinal repeated measures, SEM based analyses of genetic covariances derived from independent age restricted GWAS can be used to investigate questions regarding the stability and independence of genetic influences across time. Applying this framework to BMI across middle to late adulthood and testing two competing hypotheses, revealed that differences in BMI across age could not be explained by the accumulation of age-specific genetic influences or autoregressive processes. Instead, a common set of stable genetic influences appears to underpin genome-wide variation in BMI from middle to early old age in both men and women.

## Data Availability

All BMI and GWAS data used here are publicly available from the UK Biobank (https://www.ukbiobank.ac.uk).

## Funding/Support

This was supported in part by X grant X. REP supported by NIMH K01MH113848 and The Brain & Behavior Research Foundation NARSAD grant 28632 P&S Fund.

## Role of the Funder/Sponsor

The funding sources had no role in the preparation, review, or approval of the manuscript, or the decision to submit the manuscript for publication.

---

**Supplementary Table S1.**
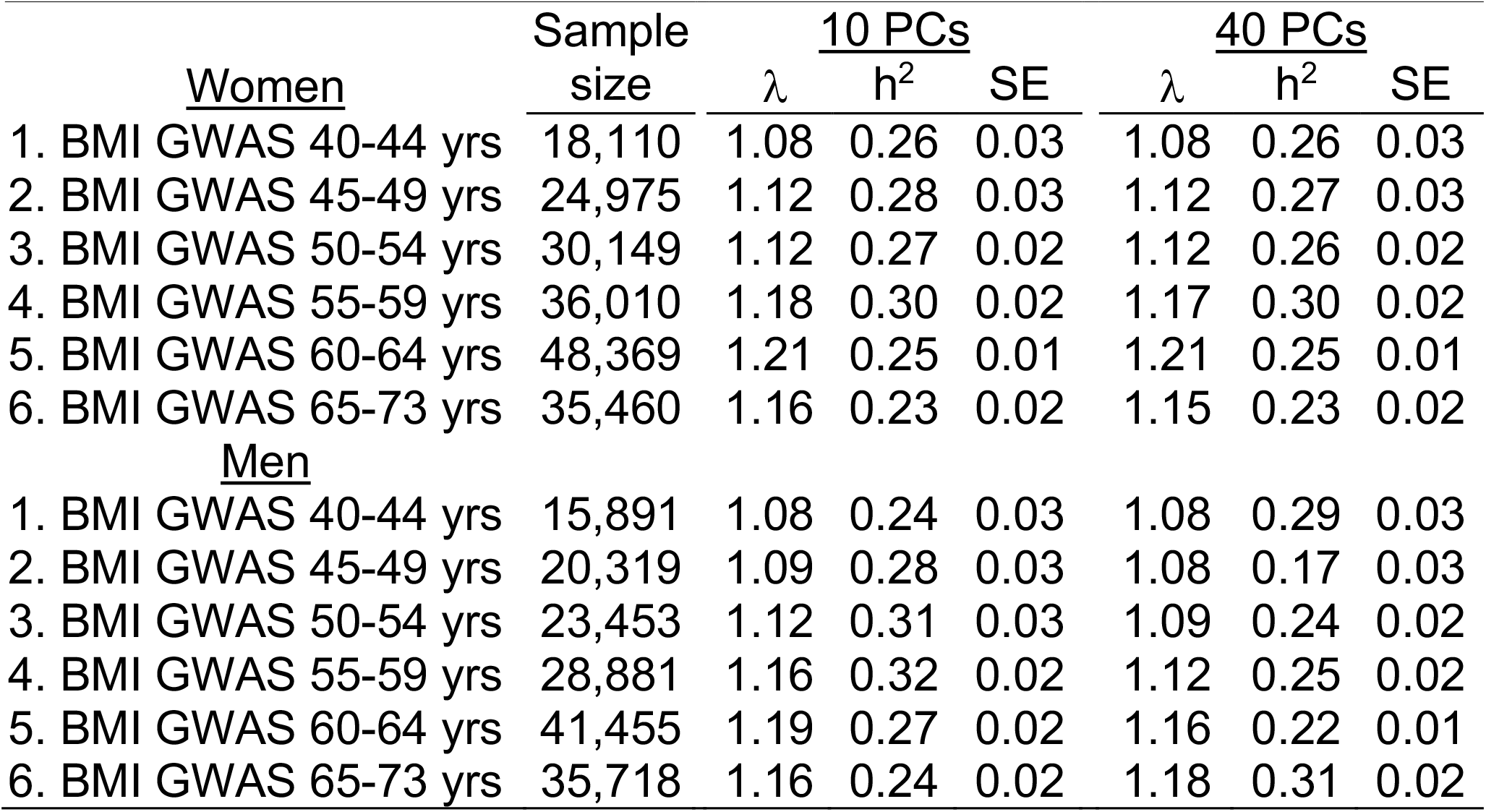
Number of men & women with complete BMI & GWAS data at each age interval, as well as genomic inflation (λ) & SNP-based heritability (h^2^) for the GWAS analyses comprising 10 versus 40 principal components (PCs).

|  | Sample size | 10 PCs |  |  | 40 PCs |  |  |
| --- | --- | --- | --- | --- | --- | --- | --- |
| | | $\lambda$ | $h^2$ | SE | $\lambda$ | $h^2$ | SE |
| <u>Women</u> |  |  |  |  |  |  |  |
| 1. BMI GWAS 40-44 yrs | 18,110 | 1.08 | 0.26 | 0.03 | 1.08 | 0.26 | 0.03 |
| 2. BMI GWAS 45-49 yrs | 24,975 | 1.12 | 0.28 | 0.03 | 1.12 | 0.27 | 0.03 |
| 3. BMI GWAS 50-54 yrs | 30,149 | 1.12 | 0.27 | 0.02 | 1.12 | 0.26 | 0.02 |
| 4. BMI GWAS 55-59 yrs | 36,010 | 1.18 | 0.30 | 0.02 | 1.17 | 0.30 | 0.02 |
| 5. BMI GWAS 60-64 yrs | 48,369 | 1.21 | 0.25 | 0.01 | 1.21 | 0.25 | 0.01 |
| 6. BMI GWAS 65-73 yrs | 35,460 | 1.16 | 0.23 | 0.02 | 1.15 | 0.23 | 0.02 |
| <u>Men</u> |  |  |  |  |  |  |  |
| 1. BMI GWAS 40-44 yrs | 15,891 | 1.08 | 0.24 | 0.03 | 1.08 | 0.29 | 0.03 |
| 2. BMI GWAS 45-49 yrs | 20,319 | 1.09 | 0.28 | 0.03 | 1.08 | 0.17 | 0.03 |
| 3. BMI GWAS 50-54 yrs | 23,453 | 1.12 | 0.31 | 0.03 | 1.09 | 0.24 | 0.02 |
| 4. BMI GWAS 55-59 yrs | 28,881 | 1.16 | 0.32 | 0.02 | 1.12 | 0.25 | 0.02 |
| 5. BMI GWAS 60-64 yrs | 41,455 | 1.19 | 0.27 | 0.02 | 1.16 | 0.22 | 0.01 |
| 6. BMI GWAS 65-73 yrs | 35,718 | 1.16 | 0.24 | 0.02 | 1.18 | 0.31 | 0.02 |

## Notes

### Competing Interest Statement

The authors have declared no competing interest.

### Summary of Updates

The middle initial of senior author Dr Roseann E. Peterson has been corrected.

## References

1 Dahl, A. K., Reynolds, C. A., Fall, T., Magnusson, P. K. & Pedersen, N. L. Multifactorial analysis of changes in body mass index across the adult life course: a study with 65 years of follow-up. International journal of obesity (2005) 38, 1133–1141, doi:10.1038/ijo.2013.204 (2014).

2 Kivimaki, M. et al. Body mass index and risk of dementia: Analysis of individual-level data from 1.3 million individuals. Alzheimers Dement 14, 601–609, doi:10.1016/j.jalz.2017.09.016 (2018).

3 Bhaskaran, K., Dos-Santos-Silva, I., Leon, D. A., Douglas, I. J. & Smeeth, L. Association of BMI with overall and cause-specific mortality: a population-based cohort study of 3.6 million adults in the UK. Lancet Diabetes Endocrinol 6, 944–953, doi:10.1016/S2213-8587(18)30288-2 (2018).

4 Nuttall, F. Q. Body Mass Index: Obesity, BMI, and Health: A Critical Review. Nutr Today 50, 117–128, doi:10.1097/NT.0000000000000092 (2015).

5 De Lorenzo, A. et al. Why primary obesity is a disease? J Transl Med 17, 169, doi:10.1186/s12967-019-1919-y (2019).

6 Ross, R. et al. Waist circumference as a vital sign in clinical practice: a Consensus Statement from the IAS and ICCR Working Group on Visceral Obesity. Nat Rev Endocrinol 16, 177–189, doi:10.1038/s41574-019-0310-7 (2020).

7 Zhang, X. H. et al. Comparison of Anthropometric and Atherogenic Indices as Screening Tools of Metabolic Syndrome in the Kazakh Adult Population in Xinjiang. International journal of environmental research and public health 13, 428, doi:10.3390/ijerph13040428 (2016).

8 Cerhan, J. R. et al. A pooled analysis of waist circumference and mortality in 650,000 adults. Mayo Clin. Proc. 89, 335–345, doi:10.1016/j.mayocp.2013.11.011 (2014).

9 Gong, J., Harris, K., Peters, S. A. E. & Woodward, M. Sex differences in the association between major cardiovascular risk factors in midlife and dementia: a cohort study using data from the UK Biobank. BMC medicine 19, 110, doi:10.1186/s12916-021-01980-z (2021).

10 Singh-Manoux, A. et al. Obesity trajectories and risk of dementia: 28 years of follow-up in the Whitehall II Study. Alzheimers Dement 14, 178–186, doi:10.1016/j.jalz.2017.06.2637 (2018).

11 Flegal, K. M., Kit, B. K., Orpana, H. & Graubard, B. I. Association of all-cause mortality with overweight and obesity using standard body mass index categories: a systematic review and meta-analysis. JAMA 309, 71–82, doi:10.1001/jama.2012.113905 (2013).

12 Parohan, M. et al. Risk factors for mortality in patients with Coronavirus disease 2019 (COVID-19) infection: a systematic review and meta-analysis of observational studies. Aging Male, 1–9, doi:10.1080/13685538.2020.1774748 (2020).

13 MacArthur, J. et al. The new NHGRI-EBI Catalog of published genome-wide association studies (GWAS Catalog). Nucleic Acids Res 45, D896–D901, doi:10.1093/nar/gkw1133 (2017).

14 Lee, S. H., Wray, N. R., Goddard, M. E. & Visscher, P. M. Estimating missing heritability for disease from genome-wide association studies. Am J Hum Genet 88, 294–305, doi:10.1016/j.ajhg.2011.02.002 (2011).

15 Yang, J. et al. Genetic variance estimation with imputed variants finds negligible missing heritability for human height and body mass index. Nat. Genet. 47, 1114–1120, doi:10.1038/ng.3390 (2015).

16 Elks, C. E. et al. Variability in the heritability of body mass index: a systematic review and meta-regression. Front Endocrinol (Lausanne) 3, 29, doi:10.3389/fendo.2012.00029 (2012).

17 Silventoinen, K. et al. Genetic and environmental factors in relative weight from birth to age 18: the Swedish young male twins study. International journal of obesity (2005) 31, 615–621, doi:10.1038/sj.ijo.0803577 (2007).

18 Cornes, B. K., Zhu, G. & Martin, N. G. Sex differences in genetic variation in weight: a longitudinal study of body mass index in adolescent twins. Behav. Genet. 37, 648–660, doi:10.1007/s10519-007-9165-0 (2007).

19 Haberstick, B. C. et al. Stable genes and changing environments: body mass index across adolescence and young adulthood. Behav. Genet. 40, 495–504, doi:10.1007/s10519-009-9327-3 (2010).

20 Stunkard, A. J., Foch, T. T. & Hrubec, Z. A twin study of human obesity. JAMA 256, 51–54 (1986).

21 Maes, H. H., Neale, M. C. & Eaves, L. J. Genetic and environmental factors in relative body weight and human adiposity. Behav. Genet. 27, 325–351, doi:10.1023/a:1025635913927 (1997).

22 Llewellyn, C. H., Trzaskowski, M., Plomin, R. & Wardle, J. From modeling to measurement: developmental trends in genetic influence on adiposity in childhood. Obesity (Silver Spring, Md.) 22, 1756–1761, doi:10.1002/oby.20756 (2014).

23 Franz, C. E. et al. Genetics of body mass stability and risk for chronic disease: a 28-year longitudinal study. Twin Res Hum Genet 10, 537–545, doi:10.1375/twin.10.4.537 (2007).

24 Silventoinen, K. & Kaprio, J. Genetics of tracking of body mass index from birth to late middle age: evidence from twin and family studies. Obes Facts 2, 196–202, doi:10.1159/000219675 (2009).

25 Cardon, L. R. in Behavior Genetic Approaches in Behavioral Medicine (eds J.R. Turner & L.R. Cardon) 133–143 (Plenum Press, 1995).

26 Bulik-Sullivan, B. K. et al. LD Score regression distinguishes confounding from polygenicity in genome-wide association studies. Nat. Genet. 47, 291–295 (2015).

27 Trzaskowski, M., Lichtenstein, P., Magnusson, P. K., Pedersen, N. L. & Plomin, R. Application of linear mixed models to study genetic stability of height and body mass index across countries and time. Int. J. Epidemiol. 45, 417–423, doi:10.1093/ije/dyv355 (2016).

28 Winkler, T. W. et al. The Influence of Age and Sex on Genetic Associations with Adult Body Size and Shape: A Large-Scale Genome-Wide Interaction Study. PLoS genetics 11, e1005378, doi:10.1371/journal.pgen.1005378 (2015).

29 Hewitt, J. K., Eaves, L. J., Neale, M. C. & Meyer, J. M. Resolving causes of developmental continuity or “tracking.” I. Longitudinal twin studies during growth. Behav. Genet. 18, 133–151 (1988).

30 Nesselroade, J. R. & Baltes, P. B. Adolescent personality development and historical change: 1970-1972. Monogr. Soc. Res. Child Dev. 39, 1–80 (1974).

31 McArdle, J. J. Latent variable growth within behavior genetic models. Behav. Genet. 16, 163–200 (1986).

32 McArdle, J. J. & Epstein, D. Latent growth curves within developmental structural equation models. Child Dev. 58, 110–133 (1987).

33 Duncan, T. E. & Duncan, S. C. A latent growth curve approach to investigating developmental dynamics and correlates of change in children’s perceptions of physical competence. Res. Q. Exerc. Sport 62, 390–398 (1991).

34 Duncan, T. E., Duncan, S. C. & Hops, H. The effects of family cohesiveness and peer encouragement on the development of adolescent alcohol use: a cohort-sequential approach to the analysis of longitudinal data. J. Stud. Alcohol 55, 588–599 (1994).

35 Hjelmborg, J. et al. Genetic influences on growth traits of BMI: a longitudinal study of adult twins. Obesity (Silver Spring, Md.) 16, 847–852, doi:10.1038/oby.2007.135 (2008).

36 Ortega-Alonso, A., Sipila, S., Kujala, U. M., Kaprio, J. & Rantanen, T. Genetic influences on adult body mass index followed over 29 years and their effects on late-life mobility: a study of twin sisters. J. Epidemiol. Community Health 63, 651–658, doi:10.1136/jech.2008.080622 (2009).

37 Ortega-Alonso, A., Sipila, S., Kujala, U. M., Kaprio, J. & Rantanen, T. Genetic influences on change in BMI from middle to old age: a 29-year follow-up study of twin sisters. Behav. Genet. 39, 154–164, doi:10.1007/s10519-008-9245-9 (2009).

38 Eaves, L. J., Long, J. & Heath, A. C. A theory of developmental change in quantitative phenotypes applied to cognitive development. Behav. Genet. 16, 143–162 (1986).

39 Boomsma, D. I. & Molenaar, P. C. The genetic analysis of repeated measures. I. Simplex models. Behav. Genet. 17, 111–123 (1987).

40 Boomsma, D. I., Martin, N. G. & Molenaar, P. C. Factor and simplex models for repeated measures: application to two psychomotor measures of alcohol sensitivity in twins. Behav. Genet. 19, 79–96 (1989).

41 Gillespie, N. A., Evans, D. E., Wright, M. M. & Martin, N. G. Genetic simplex modeling of Eysenck’s dimensions of personality in a sample of young Australian twins. Twin Res. 7, 637–648, doi:10.1375/1369052042663814 (2004).

42 Gillespie, N. A. et al. Do the genetic or environmental determinants of anxiety and depression change with age? A longitudinal study of Australian twins. Twin Res. 7, 39–53, doi:10.1375/13690520460741435 (2004).

43 Gillespie, N. A., Eaves, L. J., Maes, H. & Silberg, J. L. Testing Models for the Contributions of Genes and Environment to Developmental Change in Adolescent Depression. Behav. Genet. 45, 382–393, doi:10.1007/s10519-015-9715-9 (2015).

44 Long, E. C., Verhulst, B., Aggen, S. H., Kendler, K. S. & Gillespie, N. A. Contributions of Genes and Environment to Developmental Change in Alcohol Use. Behav. Genet. 47, 498–506, doi:10.1007/s10519-017-9858-y (2017).

45 Gillespie, N. A. et al. The genetic etiology of longitudinal measures of predicted brain ageing in a population-based sample of mid to late-age males. Neuroimage (submitted).

46 Grotzinger, A. D. et al. Genomic structural equation modelling provides insights into the multivariate genetic architecture of complex traits. Nat Hum Behav 3, 513–525, doi:10.1038/s41562-019-0566-x (2019).

47 Sudlow, C. et al. UK biobank: an open access resource for identifying the causes of a wide range of complex diseases of middle and old age. PLoS Med 12, e1001779, doi:10.1371/journal.pmed.1001779 (2015).

48 Neale, M. C. & Cardon, L. R. Methodology for Genetic Studies of Twins and Families. 1st edn, (Kluwer Academic Publishers, 1992).

49 Korkeila, M., Kaprio, J., Rissanen, A. & Koskenvuo, M. Effects of gender and age on the heritability of body mass index. Int. J. Obes. 15, 647–654. (1991).

50 UK Biobank Anthropometry, <[https://biobank.ctsu.ox.ac.uk/crystal/crystal/docs/Anthropometry.pdf> (

51 Neale, B. <https://github.com/Nealelab/> (

52 Bycroft, C. et al. The UK Biobank resource with deep phenotyping and genomic data. Nature 562, 203–209, doi:10.1038/s41586-018-0579-z (2018).

53 R Core Team. R: A language and environment for statistical computing., <https://www.R-project.org/>. (2020).

54 Rosseel, Y. Lavaan: an R package for structural equation modeling. Journal of Statistical Software 48, 1–36 (2012).

55 Lajunen, H. R. et al. Genetic and environmental effects on body mass index during adolescence: a prospective study among Finnish twins. International journal of obesity (2005) 33, 559–567, doi:10.1038/ijo.2009.51 (2009).

56 Cornes, B. K. et al. Sex-limited genome-wide linkage scan for body mass index in an unselected sample of 933 Australian twin families. Twin Res. Hum. Genet. 8, 616–632 (2005).

57 Felix, J. F. et al. Genome-wide association analysis identifies three new susceptibility loci for childhood body mass index. Hum. Mol. Genet. 25, 389–403, doi:10.1093/hmg/ddv472 (2016).

58 Couto Alves, A. et al. GWAS on longitudinal growth traits reveals different genetic factors influencing infant, child, and adult BMI. Sci Adv 5, eaaw3095, doi:10.1126/sciadv.aaw3095 (2019).

59 Sovio, U. et al. Association between common variation at the FTO locus and changes in body mass index from infancy to late childhood: the complex nature of genetic association through growth and development. PLoS genetics 7, e1001307, doi:10.1371/journal.pgen.1001307 (2011).

60 Rokholm, B. et al. Increasing genetic variance of body mass index during the Swedish obesity epidemic. PloS one 6, e27135, doi:10.1371/journal.pone.0027135 (2011).

61 Rokholm, B. et al. Increased genetic variance of BMI with a higher prevalence of obesity. PloS one 6, e20816, doi:10.1371/journal.pone.0020816 (2011).

62 Keyes, K. M. & Westreich, D. UK Biobank, big data, and the consequences of non-representativeness. Lancet 393, 1297, doi:10.1016/S0140-6736(18)33067-8 (2019).

63 Fry, A. et al. Comparison of Sociodemographic and Health-Related Characteristics of UK Biobank Participants With Those of the General Population. Am. J. Epidemiol. 186, 1026–1034, doi:10.1093/aje/kwx246 (2017).

64 Tyrrell, J. et al. Genetic predictors of participation in optional components of UK Biobank. bioRxiv (2020).

65 Silventoinen, K. et al. The CODATwins Project: The Current Status and Recent Findings of COllaborative Project of Development of Anthropometrical Measures in Twins. Twin Res Hum Genet, 1–9, doi:10.1017/thg.2019.35 (2019).

66 Silventoinen, K. et al. The CODATwins Project: The Cohort Description of Collaborative Project of Development of Anthropometrical Measures in Twins to Study Macro-Environmental Variation in Genetic and Environmental Effects on Anthropometric Traits. Twin Res Hum Genet 18, 348–360, doi:10.1017/thg.2015.29 (2015).

67 Hamer, M., Bell, J. A., Sabia, S., Batty, G. D. & Kivimaki, M. Stability of metabolically healthy obesity over 8 years: the English Longitudinal Study of Ageing. Eur. J. Endocrinol. 173, 703–708, doi:10.1530/EJE-15-0449 (2015).

68 Buscot, M. J. et al. Distinct child-to-adult body mass index trajectories are associated with different levels of adult cardiometabolic risk. Eur. Heart J. 39, 2263–2270, doi:10.1093/eurheartj/ehy161 (2018).

69 Dubois, L. et al. Genetic and environmental contributions to weight, height, and BMI from birth to 19 years of age: an international study of over 12,000 twin pairs. PloS one 7, e30153, doi:10.1371/journal.pone.0030153 (2012).

70 Eaves, L., Eysenck, H. J. & Martin, N. G. Genes, Culture, and Personality: An Empirical Approach. (Academic Press, 1989).

71 Silventoinen, K. et al. Differences in genetic and environmental variation in adult BMI by sex, age, time period, and region: an individual-based pooled analysis of 40 twin cohorts. Am. J. Clin. Nutr. 106, 457–466, doi:10.3945/ajcn.117.153643 (2017).

72 Kirkpatrick, R. M., Pritikin, J. N., Hunter, M. D. & Neale, M. C. Combining Structural-Equation Modeling with Genomic-Relatedness-Matrix Restricted Maximum Likelihood in OpenMx. Behav. Genet., doi:10.1007/s10519-020-10037-5 (2021).

73 Neale, M. C. et al. OpenMx 2.0: Extended Structural Equation and Statistical Modeling. Psychometrika 81, 535–549, doi:10.1007/s11336-014-9435-8 (2016).

